## Supplementary Figures and Tables for "Elongation Inhibitors do not Prevent the Release of Puromycylated Nascent Polypeptide Chains from Ribosomes"

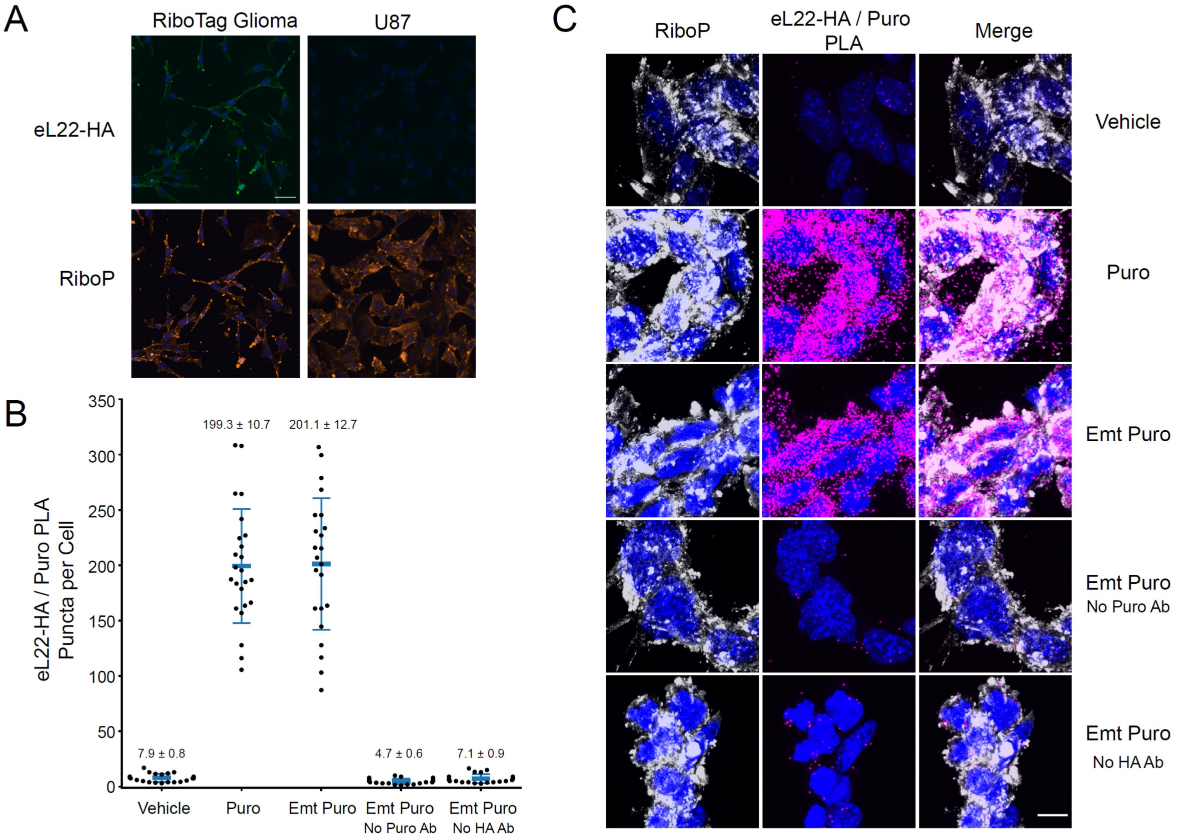

**Figure 1 – figure supplement 1: U87 glioma cells do not express eL22-HA, and eL22-HA/Puro PLA does not distinguish between emetine-treated and untreated cells at high resolution**

**(A)** Representative images (20x) of eL22-HA and RiboP immunofluorescence in RiboTag and U87 glioma cells. Scale bar, 50 µm. **(B)** Quantification of eL22-HA/Puro PLA puncta per cell. Each dot represents a cell; n=20-24 cells per condition from 2-4 separate coverslips. Blue bars represent mean and standard deviation, with mean ± SEM indicated above. **(C)** Representative confocal images (60x) of eL22-HA/Puro PLA and RiboP immunofluorescence. DAPI in blue, RiboP in white, eL22-HA/Puro PLA in magenta. Scale bar, 10 µm.

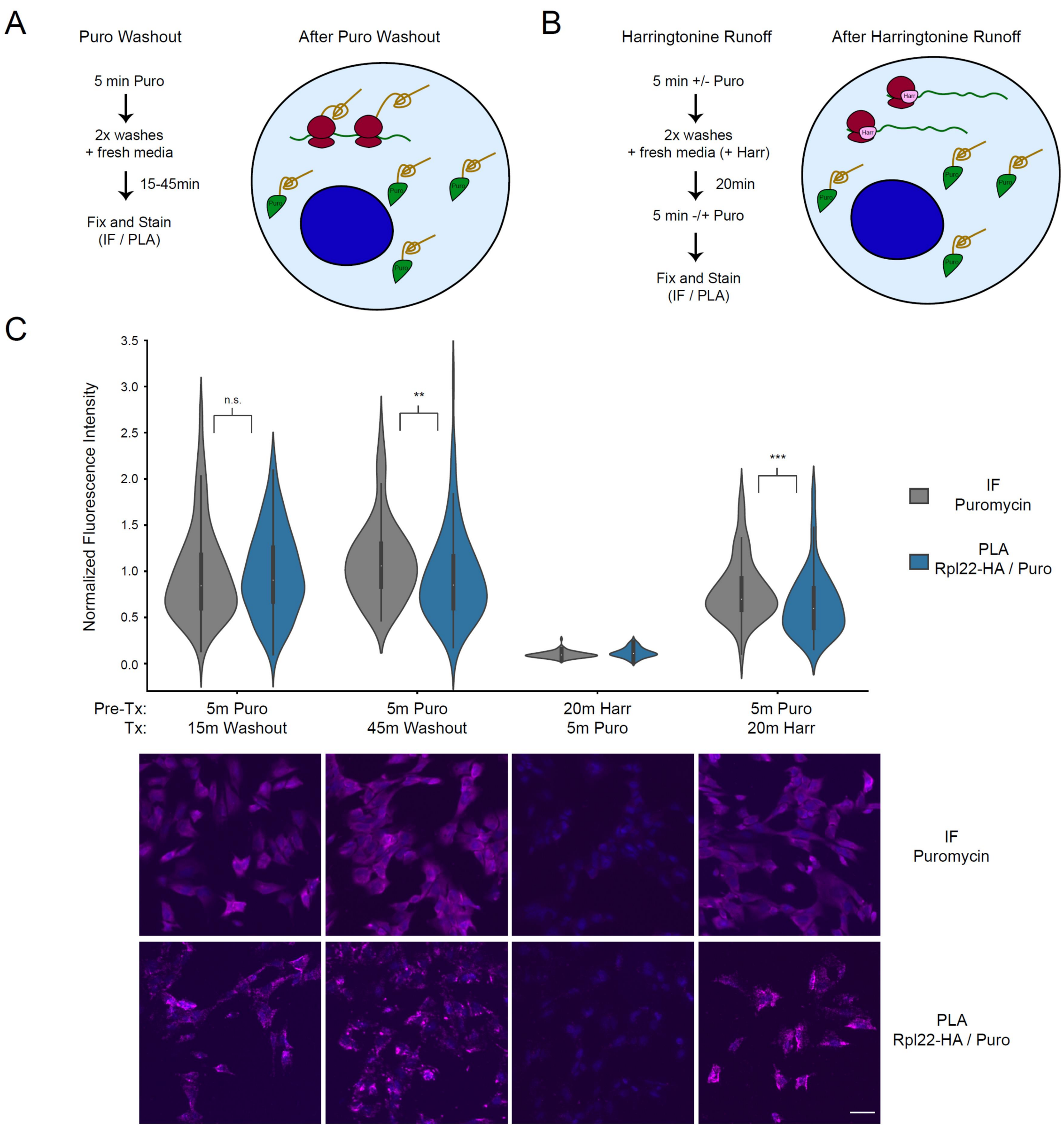

**Figure 1 – figure supplement 2: Puromycin washout and Harringtonine run-off confirm that eL22-HA/Puro PLA signal does not report on proximity of ribosomes and puromycylated nascent chains**
**(A)** Schematic depicting state of cells after puromycin washout. Cells are briefly treated with puromycin, washed, and allowed to continue in fresh media for 15-45 minutes. The vast majority of puromycylated peptides should be released from ribosomes under these conditions. **(B)** Schematic depicting state of cells after harringtonine run-off. Harringtonine stalls newly initiating ribosomes without nascent chains, while elongating ribosomes complete. The vast majority of puromycylated nascent chains should be released, and no nascent chains should be present under these conditions. **(C)** Violin plots of normalized fluorescence intensity for puromycin IF or eL22-HA/Puro PLA signal RiboTag glioma cells treated as indicated. Data are derived from 3 experiments and 96 cells per condition, and statistical comparisons indicated were conducted using Mann-Whitney U test. See **Figure 1 – figure supplement 3** for details. ** indicates $p < 0.01$, *** indicates $p < 0.001$. Representative images (10x) are shown below; DAPI in blue, puromycin IF or eL22-HA/Puro PLA in magenta. Scale bar, 50 µm.

**Figure 1 – figure supplement 3**

Statistical information pertaining to Figure 1C and Figure 1 – Supplementary Figure 2C

| Treatment | Replicates | Cells | Median | Mean | StDev | | |
|---|---|---|---|---|---|---|---|
| mRG.NoPuro_IF | 4 | 288 | 0.068 | 0.074 | 0.062 | | |
| mRG.NoPuro_PLA | 4 | 144 | 0.068 | 0.060 | 0.076 | | |
| mRG.Puro_IF | 4 | 288 | 0.930 | 0.972 | 0.433 | | |
| mRG.Puro_PLA | 4 | 144 | 0.886 | 0.924 | 0.441 | | |
| mRG.EmtPuro_IF | 4 | 288 | 0.942 | 0.969 | 0.383 | | |
| mRG.EmtPuro_PLA | 4 | 144 | 0.780 | 0.821 | 0.408 | | |
| mRG.EmtPuroNoHA_IF | 3 | 144 | 0.862 | 0.883 | 0.357 | | |
| mRG.EmtPuroNoHA_PLA | 3 | 112 | 0.052 | 0.055 | 0.048 | | |
| mRG.EmtPuroNoPuro_IF | 3 | 144 | 0.073 | 0.074 | 0.024 | | |
| mRG.EmtPuroNoPuro_PLA | 3 | 112 | 0.099 | 0.097 | 0.084 | | |
| mRG.AnisoPuro_IF | 4 | 288 | 0.122 | 0.136 | 0.107 | | |
| mRG.AnisoPuro_PLA | 3 | 112 | 0.057 | 0.055 | 0.059 | | |
| U87.EmtPuro_IF | 3 | 264 | 0.844 | 0.875 | 0.388 | | |
| U87.EmtPuro_PLA | 3 | 112 | 0.150 | 0.162 | 0.138 | | |
| mRG.PuroWash15_IF | 3 | 128 | 0.906 | 0.970 | 0.510 | | |
| mRG.PuroWash15_PLA | 3 | 128 | 0.942 | 0.980 | 0.460 | | |
| mRG.PuroWash45_IF | 3 | 96 | 1.081 | 1.105 | 0.436 | | |
| mRG.PuroWash45_PLA | 3 | 96 | 0.910 | 0.970 | 0.569 | | |
| mRG.HarrPuro_IF | 3 | 96 | 0.095 | 0.096 | 0.035 | | |
| mRG.HarrPuro_PLA | 3 | 96 | 0.114 | 0.118 | 0.052 | | |
| mRG.PuroWashHarr_IF | 3 | 96 | 0.746 | 0.795 | 0.333 | | |
| mRG.PuroWashHarr_PLA | 3 | 96 | 0.619 | 0.640 | 0.342 | | |
| **Group1** | **Group2** | **Median Group1** | **Median Group2** | **U** | **p** | **$n_1$** | **$n_2$** |
| mRG.Puro_IF | mRG.EmtPuro_IF | 0.930 | 0.942 | 41096 | 0.4254 | 288 | 288 |
| mRG.Puro_PLA | mRG.EmtPuro_PLA | 0.886 | 0.780 | 8739 | 0.0106 | 144 | 144 |
| mRG.Puro_IF | mRG.Puro_PLA | 0.930 | 0.886 | 18997 | 0.0776 | 288 | 144 |
| mRG.EmtPuro_IF | mRG.EmtPuro_PLA | 0.942 | 0.780 | 15544 | 1.10E-05 | 288 | 144 |
| mRG.EmtPuroNoHA_IF | mRG.EmtPuroNoHA_PLA | 0.862 | 0.052 | 15 | 5.47E-43 | 144 | 112 |
| U87.EmtPuro_IF | U87.EmtPuro_PLA | 0.844 | 0.150 | 454 | 2.67E-50 | 264 | 112 |
| mRG.PuroWash15_IF | mRG.PuroWash15_PLA | 0.906 | 0.942 | 7736 | 0.220959 | 128 | 128 |
| mRG.PuroWash45_IF | mRG.PuroWash45_PLA | 1.081 | 0.910 | 3442 | 0.001233 | 96 | 96 |
| mRG.PuroWashHarr_IF | mRG.PuroWashHarr_PLA | 0.746 | 0.619 | 3200 | 1.28E-04 | 96 | 96 |

Summary of replicates, cells, and statistical testing (Mann-Whitney U) for indicated treatment groups.

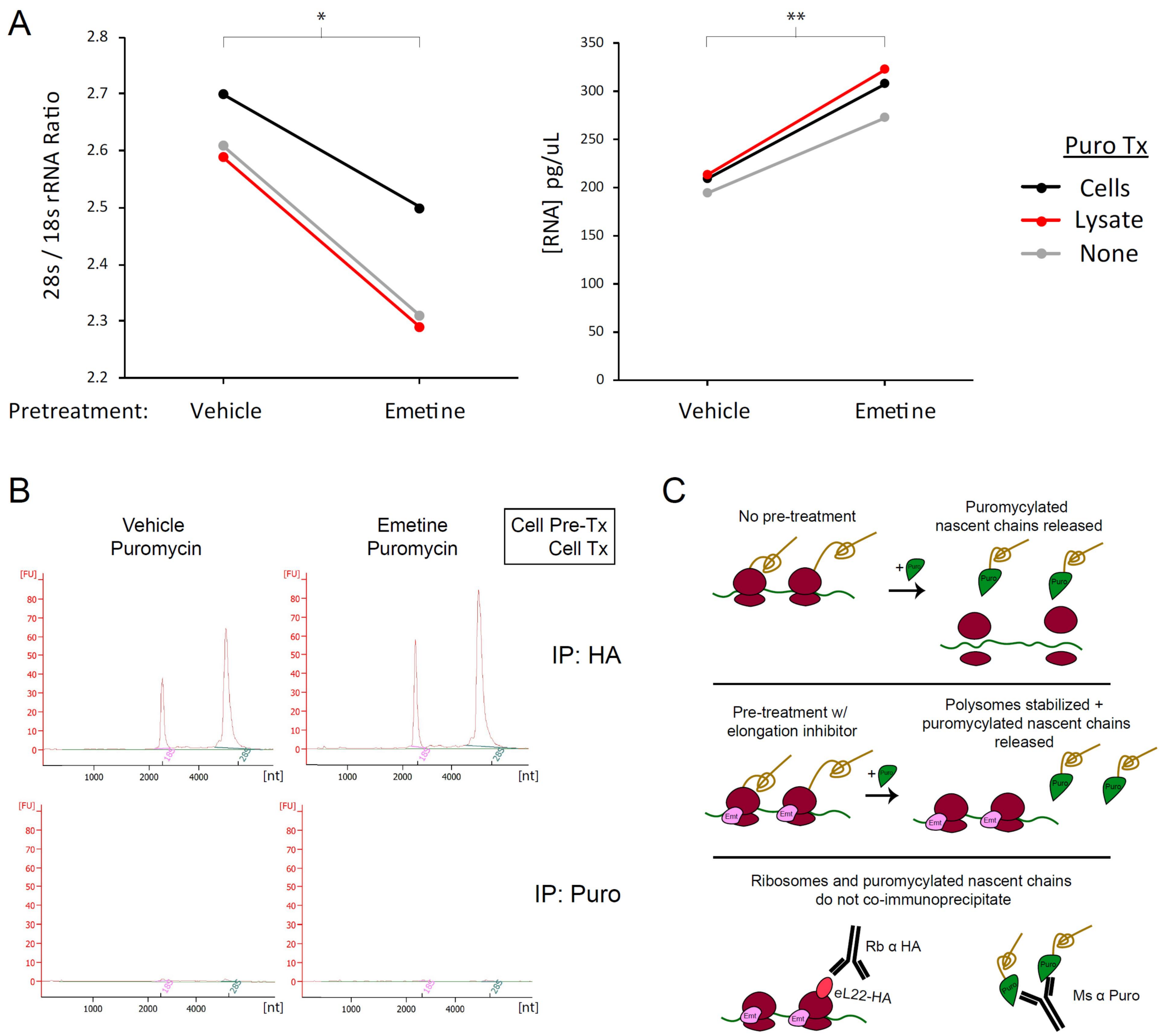

**Figure 2 – figure supplement 1: Pretreatment with emetine increases yield of intact ribosomes in eL22-HA immunoprecipitation, but does not enable capture of ribosomal RNA in puromycin immunoprecipitation.**

**(A)** Regardless of puromycin treatment condition, emetine increases intact ribosome capture as shown by lower 28s/18s rRNA ratio (*left*; IP is against eL22-HA and therefore captures more free large subunits in the absence of emetine) and higher total RNA yield (*right*). * indicates $p < 0.05$, ** indicates $p < 0.01$ for paired t-test. *Le*ft: $t(2) = 8.000$, $p = 0.0153$. *Right*: $t(2) = 10.580$, $p = 0.0088$. **(B)** RNA Pico Bioanalyzer of total RNA eluted from immunoprecipitation of eL22-HA or puromycin. **(C)** Schematic summarizing results of co-immunoprecipitation experiments.

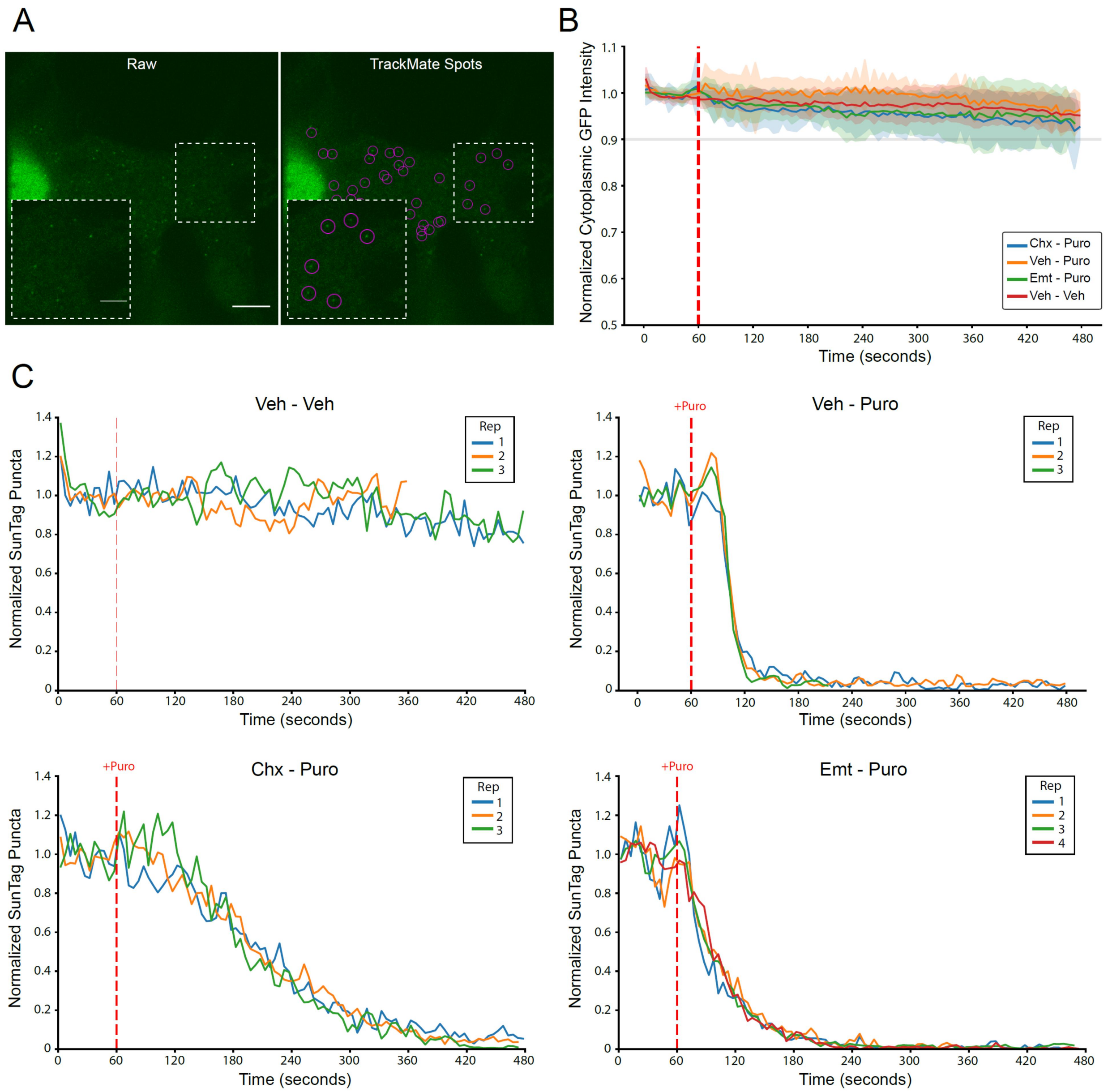

**Figure 3 – figure supplement 1: SunTag puncta detection, minimal photobleaching, and consistency of replicate imaging trials**

**(A)** Representative images of SunTag puncta quantification using TrackMate spot detection; registered spots are circled in purple. Scale bars: 10 µm for large field, 5 µm for inset. **(B)** Live imaging time course of background cytoplasmic GFP intensity from cells treated as indicated. Cytoplasmic ROIs for each cell were normalized to the average GFP intensity in that cell during the initial 10 frames. Data correspond to the exact cells presented in **Figure 3C**. Puromycin was added at 60 seconds into the 8-minute imaging trial (dashed red line). Plotted are the mean ± standard deviation of all cells, computed in five second time intervals. **(C)** Live imaging time course of normalized SunTag puncta for the exact cells presented in **Figure 3C**. Plotted is the mean of all cells (n=7-12) in each replicate imaging trial for each treatment condition as indicated. Puromycin was added at 60 seconds into the 8-minute imaging trial (dashed red line).

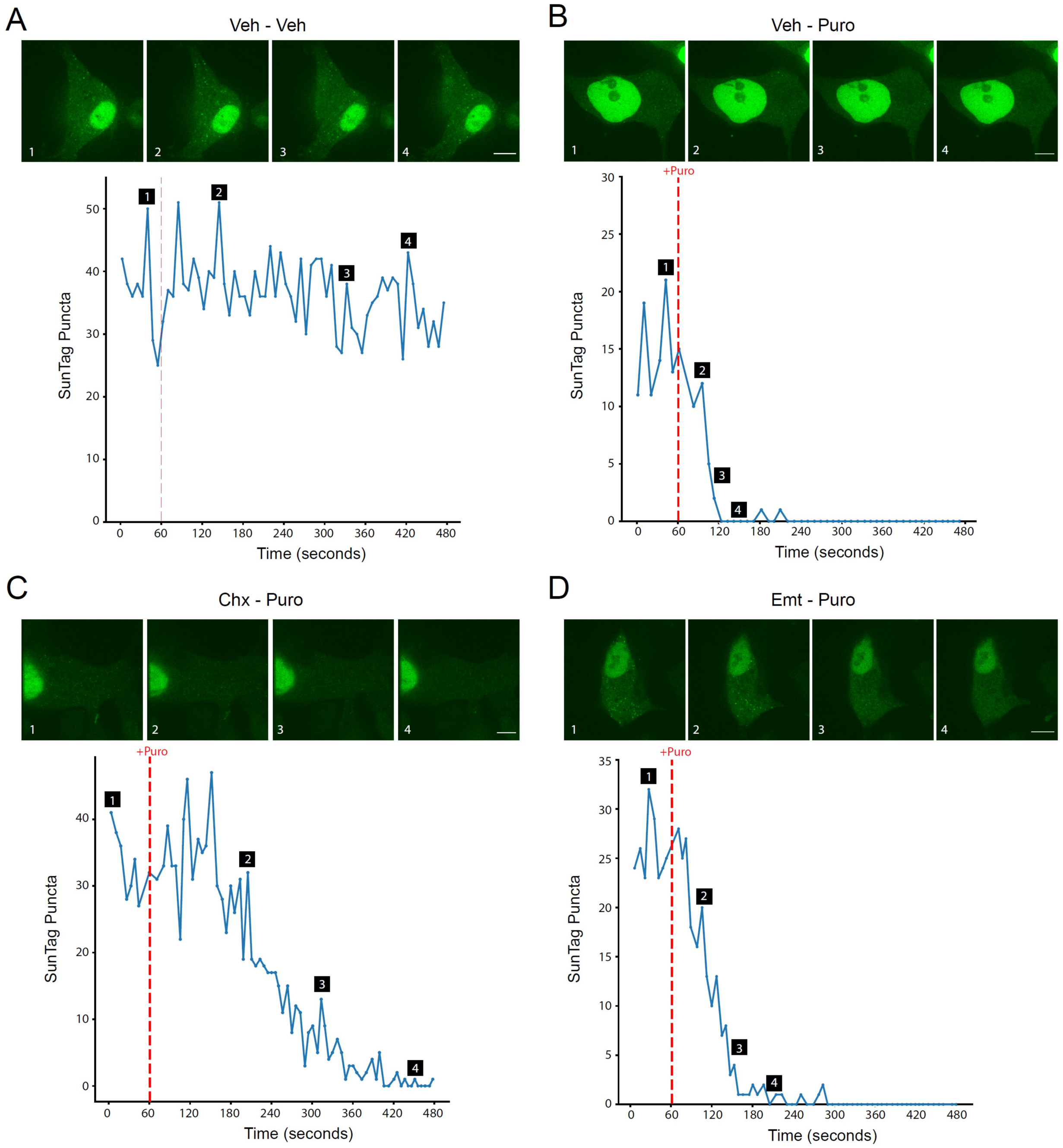

**Figure 3 – figure supplement 2: Representative images and traces of single cell SunTag imaging**
**(A-D)** Representative single cell raw puncta counts from each frame are plotted during live imaging with treatments as indicated. Puromycin was added at 60 seconds into the 8-minute imaging trial (dashed red line). Images of cells are marked with labels 1-4, corresponding to the same labels on the time course of raw puncta counts plotted below. Scale bars: 10 µm.

**Figure 3 – video 1: Time lapse live imaging of puromycin-induced SunTag puncta disappearance**
Representative time lapse imaging of SunTag puncta disappearing upon puromycin treatment, corresponding to **Figure 3C**. Cycloheximide or emetine were added 4 minutes before the start of imaging (5-minute pre-treatment), and puromycin was added 60 seconds into the 8-minute imaging trial (red Puro text indicates when Puro is added in each trial). Frames for each trial were upscaled to the lowest common denominator, and subsequently down-sampled to 102 frames, which are streamed at 8 frames per second (see Methods). Playback is approximately ~40x (~8 minutes in ~12 seconds). The exact timestamp of each image in each trial is indicated. Scale bar: 10 µm.

**Figure 4 – figure supplement 1**

Summary of MolProbity analysis of IgG2a Fab alignments within 80S A site

| Model # | MolProbity Clashscore[a] | Structures of Comparable Resolution With Worse MolProbity Clashscores (%)[b] |
|---|---|---|
| 1 | 246.30 | 0 |
| 2 | 196.19 | 0 |
| 3 | 183.82 | 0 |
| 4 | 177.55 | 0 |
| 5 | 155.58 | 0 |
| 6 | 143.09 | 0 |
| 7 | 134.19 | 0 |
| 8 | 126.94 | 0 |
| 9 | 107.07 | 0 |
| 10 | 105.83 | 0 |
| 11 | 91.18 | 0 |
| 12 | 85.79 | 0 |
| Best Fit in A Site | 22.89 | 25 |

| Structure | PDB ID | MolProbity Clashscore[a] | Structures of Comparable Resolution With Worse MolProbity Clashscores (%) |
|---|---|---|---|
| Ms IgG2a | 1IGT | 20.63 | 87[c] |
| Rb 80S | 6SGC | 7 | 87[b] |

[a] The number of serious steric overlaps (> 0.4 Å) per 1000 atoms

[b] Out of 1,784 structures solved using cryo-EM at any resolution

[c] Out of 141 structures solved with X-ray crystallography with resolutions of 2.80 ± 0.25 Å
